# Macroevolutionary drivers of morphological disparity in the avian quadrate

**DOI:** 10.1101/2023.10.05.561045

**Authors:** Pei-Chen Kuo, Guillermo Navalón, Roger B. J. Benson, Daniel J. Field

## Abstract

In birds, the quadrate bone acts as a hinge between the lower jaw and the skull, playing an important role in cranial kinesis. As such, the evolution of avian quadrate morphology may plausibly be assumed to have been influenced by selective pressures related to feeding ecology. However, variation in quadrate morphology across living birds and its potential relationship with ecology have never been quantitatively characterised. Here, we used three-dimensional geometric morphometrics and phylogenetic comparative methods to quantify morphological variation of the quadrate and its relationship with an array of key ecological features across 200 bird species covering all major extant lineages. We found non-significant associations between quadrate shape and several aspects of feeding and foraging ecology across different scales of phylogenetic comparison. By contrast, allometry and phylogeny exhibit stronger relationships with quadrate shape than other ecological features. We show that the avian quadrate evolves as an integrated unit and exhibits strong associations with the morphologies of neighbouring bones. Our results collectively suggest a macroevolutionary scenario in which the shape of the quadrate evolved jointly with other elements of the avian kinetic system, with the major lineages of birds exploring alternative quadrate morphologies—highlighting the potential diagnostic value of quadrate morphology in investigations of fossil bird systematics.

## 1. Introduction

Crown birds (Neornithes) are the most species-rich group of extant tetrapods, showcasing an extraordinary diversity of body forms and ecological disparity across virtually all major subaerial environments [1, 2]. The origin of much of this diversity is thought to lie in a large-scale adaptive radiation in the aftermath of the K-Pg mass extinction [e.g., 3]. This episode of diversification rapidly gave rise to ecologically disparate avian lineages [4] and was characterised by the rapid accumulation of morphological disparity across the skeleton [e.g., 5], including striking variability in the morphology and ecological specialisation of the feeding apparatus [e.g., 6-7]. Recent research has challenged assumptions regarding straightforward relationships between dietary traits and large-scale macroevolutionary patterns of craniofacial shape variation (e.g., beak shape [8]; cranial shape [9]; but see [10]) although more nuanced relationships may exist at more restricted phylogenetic scales (e.g., in anseriforms, [11]; and charadriiforms, [12]; although see [13]) and/or with finer anatomical traits within specific regions of the head [14].

Despite the biomechanical importance of the palate system for feeding in neornithine birds [e.g., 6, 15], detailed investigations of the factors underlying the evolution of avian palatal disparity are scarce (although see [16] exploring allometry of the vomer). Within this system, the quadrate bone (*os quadratum* [17], hereafter ‘quadrate’) serves a key role in feeding biomechanics and cranial kinesis as it articulates directly with numerous components of the feeding apparatus: the mandible, the bony palate via the pterygoid bone, and the rostrum via the quadratojugal’s connection with the jugal bars [18–21] (Fig. 1). During jaw closure, the quadrate is pulled rostrally by the action of the pterygoideus muscle group and transfers adduction forces through two set of pushrods (the pterygoid-palatine complex and the jugal bars) to the mandible and rostrum which ultimately causes dorsiflexion of the beak at kinetic positions situated at its base and/or at its tip [22]. Furthermore, the quadrate provides attachment sites for some key muscles, such as *M. protractor pterygoidei et quadrati*, *M. pseudotemporalis profundus*, and *M. adductor mandibulae caudalis*, which are involved in the movement of both the upper and lower jaws [19]. Different neornithine lineages have modified this basic configuration into a plethora of disparate morphologies thought to be tuned to functions and behaviours related to procuring different types of food [6]. As such, the considerable morphological variation exhibited by avian quadrates may reflect this ecological and functional variation. However, since the quadrate is part of a functionally integrated kinetic system involving several interconnected skeletal components, macroevolutionary patterns in quadrate geometric disparity may also reflect constraints imposed by the morphological evolution of its neighbouring bones.

**Fig. 1.**
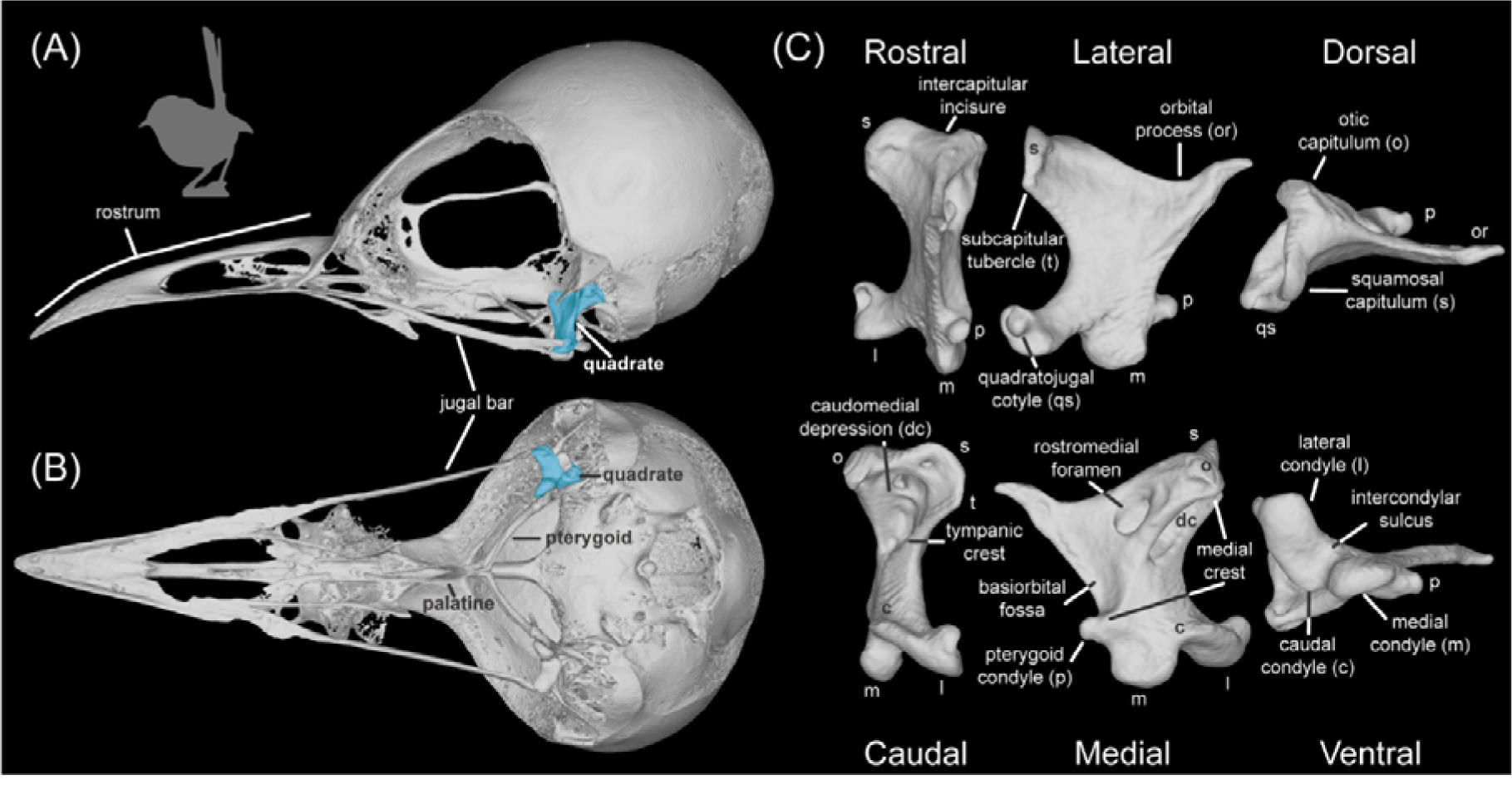
Quadrate anatomy in Neornithes. The skull of *Malurus melanocephalus* (UMMZ 224775), the species exhibiting a quadrate shape closest to the mean shape in our dataset in lateral (A) and ventral view (B), and its quadrate in rostral, lateral dorsal, caudal, medial, and ventral views (C). The quadrate of *Malurus melanocephalus* is highlighted in blue in (A) and (B).

Here, we quantitatively explore macroevolutionary links between the morphology of the avian quadrate and feeding, foraging and habitat ecology across a wide range of extant birds, while taking into consideration functional and developmental relationships of this crucial skeletal element with other osteological components of the palate and skull. Our work clarifies the extent to which adaptation for feeding function has driven morphological disparity of a key component of the avian feeding apparatus and highlights the influence of structural constraints and morphological integration on dictating skeletal geometry.

## 2. Materials & Methods

### (a) Dataset, phylogenetic hypothesis and ecological information

Our dataset includes 200 bird species and samples most major neornithine subclades (Fig 2; Supplementary Materials). A time-calibrated phylogeny of these 200 species was constructed using the backbone of a recent fossil-calibrated genome-level molecular analysis [4] and several other subclade-focused studies for the interrelationships within the following neornithine clades: Tinamiformes [23], Galliformes [24], Anseriformes [25], Apodiformes [26], Rallidae [27], Podicipedidae [28], Accipitridae [29], Psittaciformes [30], and Passeriformes [31]. Our topology was constructed using Mesquite v. 3.61 [32], and the resulting tree was temporally calibrated using *paleotree* v.3.4.4 [33] within the *R* statistical environment v. 4.2.1 [34]. Downstream analyses were also undertaken in *R*. We sourced divergence time estimates from recent genome-level molecular analyses [4, 31], and the ages of remaining uncalibrated nodes lacking recent molecular divergence time estimates were estimated using the “timePaleoPhy” function of *paleotree*, using the ‘equal’ time-scaling method.

**Fig. 2.**
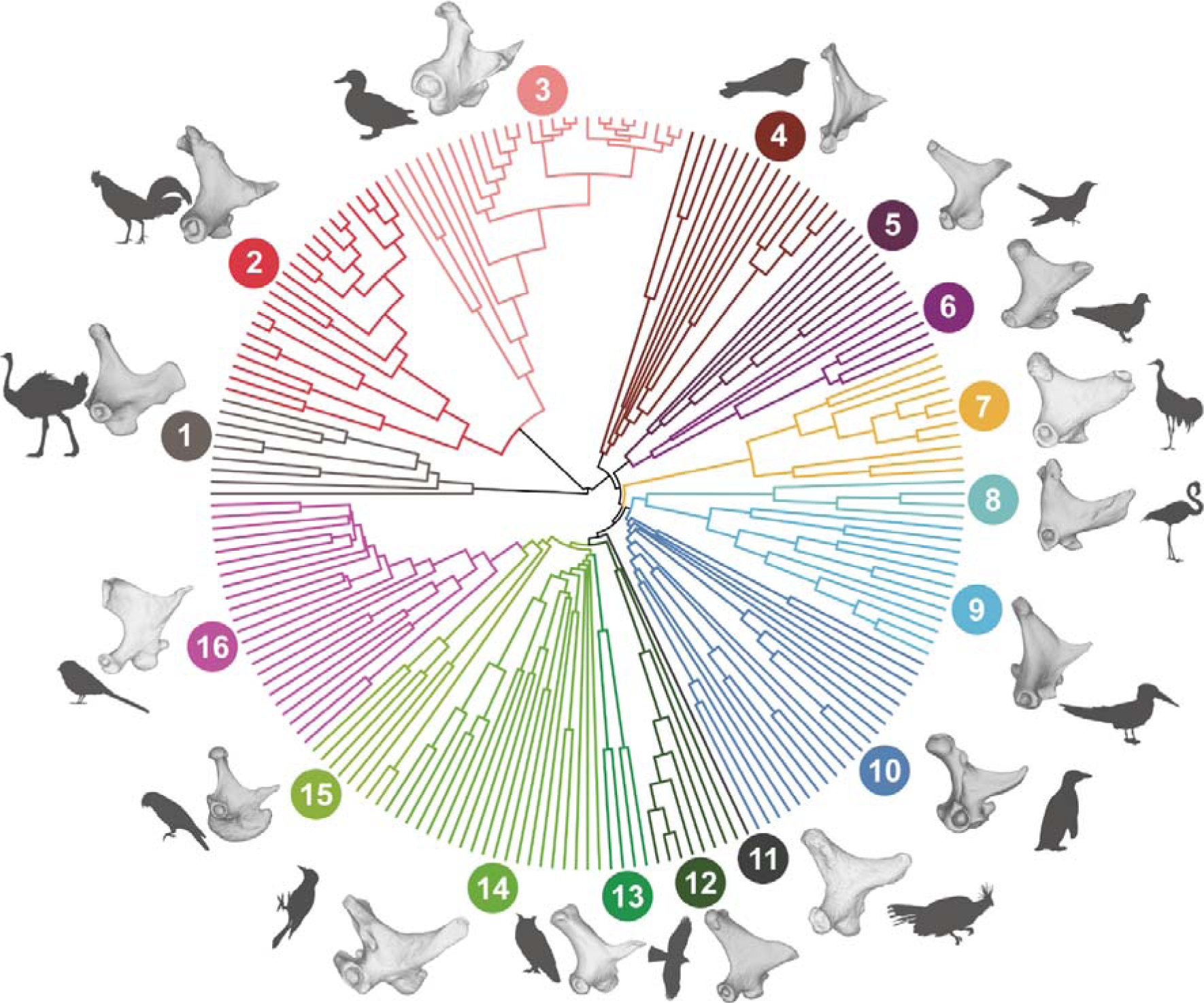
Quadrate geometric disparity in Neornithes. Time-calibrated phylogeny of the taxa included in this study (n = 200) including exemplar quadrate morphologies for each clade. 1—Palaeognathae (*Struthio camelus* UMZC uncatalogued), 2—Galliformes (*Alectura lathami* NHMUK S/2010.1.31), 3—Anseriformes (*Anas aucklandica* NHMUK S 2006.39.1), 4—Strisores (*Eurostopodus mysticalis* NHMUK S 1981.95.6), 5—Otidimorphae (*Hierococcyx fugax* FMNH 357420), 6—Columbimorphae (*Treron capellei* UMMZ 220454), 7—Gruiformes (*Balearica pavonina* NHMUK 1859.10.26), 8—Mirandornithes (*Phoenicopterus roseus* UMZC 346.B), 9—Charadriiformes (*Rynchops niger* FMNH 376309), 10—Ardeae (*Spheniscus humboldti* NHMUK S 2000.7), 11—Opisthocomiformes (*Opisthocomus hoazin* NHMUK 1961.6.1), 12—Accipitriformes (*Accipiter nisus* NHMUK S 1982.149.1), 13—Strigiformes (*Tyto alba* NHMUK S 1899.22.1), 14— Coraciimorphae (*Picus viridis* NHMUK S 1982.5.1), 15—non-passerine Australaves (*Nestor notabilis* FMNH 23530), 16—Passeriformes (*Malurus melanocephalus* UMMZ 224775).

Feeding and foraging autecology of each species in our dataset was characterised using eight classes of ecological information: trophic level (discrete: herbivore, carnivore, scavenger), trophic niche (discrete: frugivore, granivore, nectarivore, terrestrial herbivore, aquatic herbivore, invertivore, vertivore, aquatic predator, scavenger), habitat (discrete: forest, woodland, riverine, grassland, wetland, shrubland, human modified, marine, coastal, and rock), primary foraging lifestyle (discrete: aerial, insessorial, terrestrial, aquatic and generalist), and main diet (two categories: diet and diet.50). Diet was treated as discrete following the EltonTraits 1.0 database [35] and was divided into five categories: fruinect, invertebrate, omnivore, plantseed and vertfishscav. Diet.50 was also treated as discrete (fruit, invertebrate, nect, omnivore, planto, plantseed, seed, vert.ect, vert.end, vert.fish and vertfishscav). The Diet.50 variable is based on the semiquantitative dietary preference data from the EltonTraits 1.0 database and these semiquantitative dietary preferences were composed of ten different dietary categories coded as 10 binned increments representing the importance of each item in the diet of each species. Main diet (Diet.50) follows the majority dietary preference (> 50%); for taxa whose majority dietary preference is lower than 50%, the first two major dietary preferences were chosen. Additional variables examined were quadrate size (continuous, quantified as log_10_-transformed centroid size, see below), semiquantitative dietary preferences (binned in 10 increments, see above) and the use of the beak during feeding (discrete; cracking/ripping, tearing, pecking/grazing, grabbing/gleaning, probing, filtering). Variation by taxonomic was also assessed. Data on trophic level, trophic niche, habitat, and primary foraging lifestyle were retrieved from the global AVONET database [36], and the size of the quadrate was estimated from the centroid size of landmark constellations as one of the outputs of the “gpagen” function of the *geomorph* 4.04 *R* package [37]. Semiquantitative dietary preferences for each species were obtained from the global EltonTraits 1.0 database, and this noncontinuous matrix was transformed into a Euclidean distance matrix using the function “dist” from the *R* package *stats* v.3.6.2 [34], which we then transformed using a principal coordinates analysis (PCoA). The scores from this PCoA were then used as variables (diet.matrix) for downstream analyses. To code the use of the beak during feeding (UBF) we followed the methods described in [8].

### (b) Quadrate shape

We captured the complex morphology of the quadrate bone from each species by means of landmark-based geometric morphometrics applied to three-dimensional surfaces extracted from micro-CT scanned museum specimens. Specimens came from three sources: 1) CT scans of *Ptilopachus petrosus* were obtained from the Natural History Museum, Tring; 2) Numerous specimens housed in the University of Cambridge Museum of Zoology were scanned specifically for this study; and 3) remaining scans from several collections around the world were sourced from the project TEMPO Birds, managed by R.B.J.B., downloaded from the online repository MorphoSource (https://www.morphosource.org/; Supplementary Materials). Three-dimensional volumes were generated from VGSTUDIO MAX 3.5.4 (Volume Graphics) and Avizo 2019.3 (Thermo Fisher Scientific). After segmenting each quadrate, internal voids were filled in Avizo to provide a solid surface and prevent artefacts during the sliding treatment of curve and surface semilandmarks. Finally, three-dimensional volumes were transformed as three-dimensional surfaces for landmarking in Avizo. Our landmarking scheme is a modified version of [38] and was focused on characterising: (1) the articular surfaces between the quadrate and its neighbouring bones; (2) the position of muscle attachments; and (3) the overall shape of the quadrate (a detailed description of our landmarking scheme, including anatomical criteria used to place our shape coordinates, is presented in the Supplementary Materials). Series of curve semi-landmarks and patches of surface semi-landmarks were initially positioned using arbitrary numbers of points for each specimen, and then resampled to equal counts of evenly-spaced semi-landmarks before downstream analysis using the “digit.curves” function of *geomorph* (see [39]; for further details see Supplementary Materials). The final densely sampled landmark configurations (eight landmarks, 509 curve semi-landmarks, and 797 surface semi-landmarks) were subjected to a Generalized Procrustes Analysis to remove geometric information related to scaling, translation and rotation using the “gpagen” function in *geomorph*. The Minimum Bending Energy sliding method [39, 40] was used to slide the majority of semilandmark coordinates (see Supplementary Materials for details about digitizing surface semilandmarks and Generalized Procrustes Analysis).

### (c) Principal Components Analysis

Shape data (Procrustes coordinates) were subjected to a Principal Components Analysis (PCA) to visualize quadrate shape variation using the “plotTangentSpace” function in *geomorph*. Shape changes associated with each of the first five major axes of total quadrate shape variation (PC1-PC5) were displayed as deformations warped on the three-dimensional surface of the quadrate of the individual species closest to the mean quadrate shape in our sample, which was Red-backed Fairywren, *Malurus melanocephalus*. Specifically, this three-dimensional surface and the mean shape from the sample were projected onto the scores representing the 0.05 and 0.95 quantiles for each PC axis by means of thin-plate spline deformation [41] using the function “tps3d” from the package *Morpho* v. 2.10 [42] and “shape.predictor” from *geomorph*. We also plotted the respective landmark configurations onto the deformed meshes using “shape.predictor”, and coloured landmark constellations according to per-landmark-variances using the ‘hot.dots’ function (freely available following this link: https://zenodo.org/record/3929193).

### (d) Multivariate statistics

We ran Phylogenetic Generalized Least Squares (PGLS) linear models of differing complexity (see Supplementary Materials: Additional results – PGLS) to evaluate the main effects and interactions of all the ecological variables, size (log_10_ centroid size) and phylogenetic group with respect to quadrate shape variance using the “procD.pgls” function in *geomorph*. Since this function assumes a single-rate Brownian motion model of evolution [37], we re-scaled the branches of our calibrated phylogenetic tree to adjust the phylogenetic-variation-covariation matrix to a Brownian motion process (e.g., [43]). For this, we ran each PGLS regressions once and then estimated the value of Pagel’s lambda in the residuals from the PGLS regression using the “phylosig” function from the package *phytools* v.1.03 [44] (for all values of lambda see Supplementary Materials). Then, we used this value of Pagel’s lambda to rescale our phylogeny using the function “rescale” in the package *geiger* v. 2.0.10 [45] and ran each PGLS regression a second time.

Finally, we explored shape covariation between quadrate geometry and the shapes of neighbouring bones (i.e. evolutionary integration; see Supplementary tables S5-S8 and Figure 5 for the full list of relationships tested). To do so we obtained shape data for the pterygoid-palatine complex (PPC, vomer excluded), jugal, cranium, rostrum, and mandible from [46] and pruned our dataset to match the species list from that study. The pruned quadrate dataset (n = 97) was subjected to Phylogenetic Two Blocks Partial Least Squares (p-2BPLS) using the function “phylo.integration” in *geomorph* and a rescaled phylogeny using the lambda value estimated from shape residuals from the simplest PGLS linear model (shape ∼ 1) to comply with the assumption of Brownian Motion from p-2BPLS. We conducted p-2BPLS among a number of variables, including: 1) shape (Procrustes coordinates) of the quadrate and the shape of each bone with which it articulates; 2) shape of the individual regions of the quadrate (i.e. internal integration); and 3) shape of the individual regions of the quadrate and the shape of each bone with which the quadrate articulates. The full list of variables explored is available in the Supplementary Materials. Because different relationships between morphology, ecology and integration can emerge at different phylogenetic scales [8, 9], we repeated all the above mentioned analyses across Neornithes as a whole, as well as subsets corresponding to Telluraves (landbirds), and non-Telluraves.

## 3. Results

### (a) Main patterns of quadrate shape variation in Neornithes

The first five principal components (PC) account for only 59.82% of total avian quadrate shape variance (PC1: 28.30%; PC2: 11.64%; PC3: 7.66%; PC4: 6.54%; PC5: 5.48%; see Fig. S12 and see Fig. S16 for details of shape patterns associated with each major axis), and each subsequent PC explains less than 5% of total shape variance.

PC1 separates Galloanserae (Galliformes + Anseriformes, with positive scores) from other Neornithes (mostly with negative scores, see the dark grey convex hull in Fig. 3 and Fig. S13 for a version of this figure coloured by clade). This PC axis describes important differences in the otic process, the quadratojugal cotyle, the pterygoid condyle, the mandibular process, and the quadrate body itself (Figs. 1, 3 and S16). Quadrates associated with a narrow otic process (i.e. with the two otic capitula positioned close to each other), a narrow mandibular process (i.e. the medial and lateral condyle are rostrocaudally aligned), and a slender quadrate body, such as those of Galloanserae, occupy the positive side of PC1 (Figs. 3; S13, A-D, PC1 positive). On the other hand, quadrates with negative scores on PC1 display a wider otic process (the otic capitula are relatively widely separated), a laterally positioned quadratojugal cotyle, a rostrally oriented pterygoid condyle, a caudally expanded mandibular process (i.e. the caudal condyle is well-developed) and a wider quadrate body (Figs. 3 and S16, PC1 negative).

**Fig. 3.**
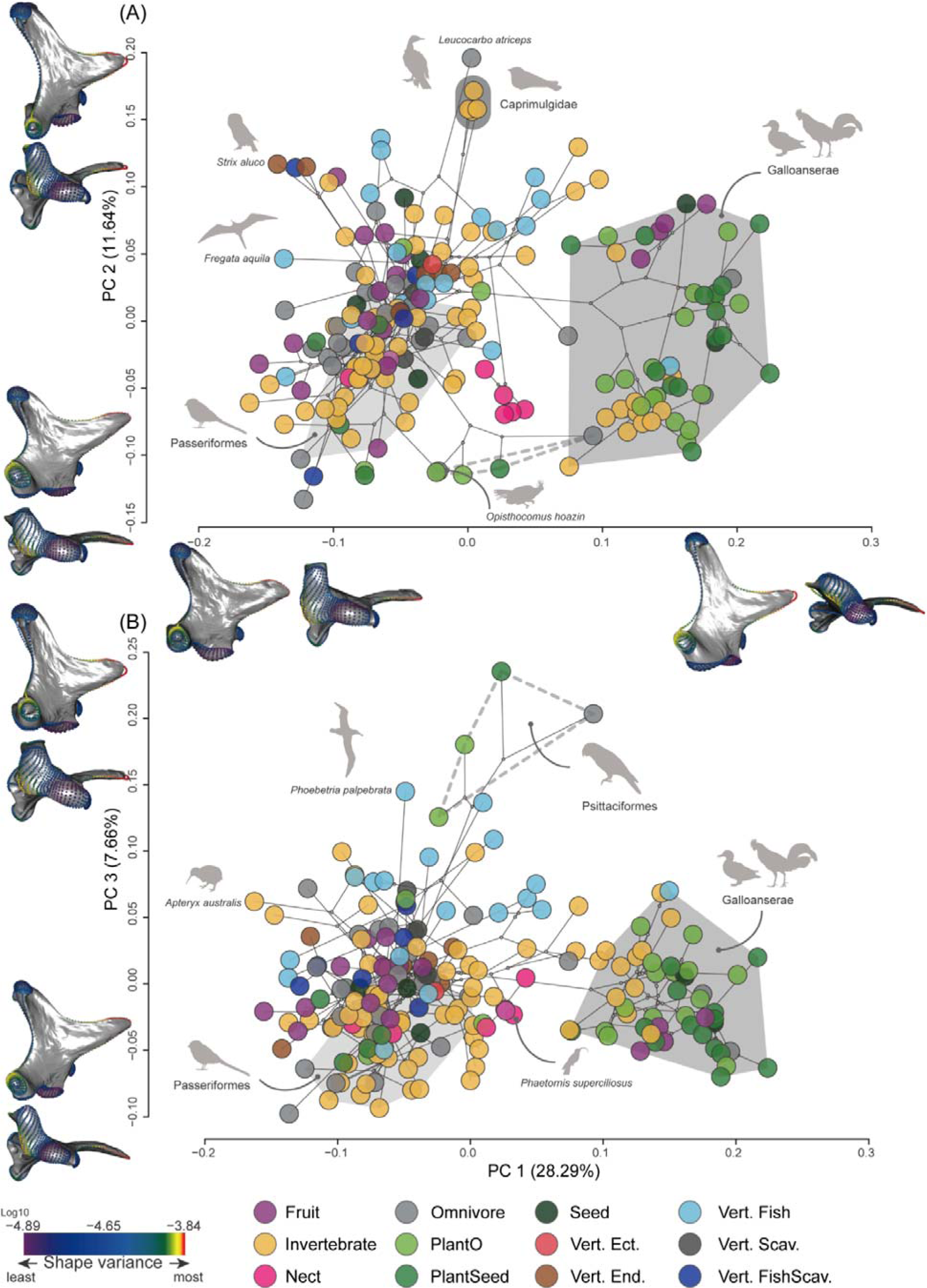
Macroevolutionary patterns of quadrate shape variation and dietary ecology in Neornithes. Phylomorphospace of the first three axes of quadrate shape variation. (A) PC1-PC2 and (B) PC1-PC3. Quadrate models are shown in lateral and ventral views and represent 0.95 quantiles of shape variance along PC2 and PC3 (top) in (A) and (B), respectively; 0.05 quantiles of shape variance along PC2 and PC3 (bottom) in (A) and (B), respectively; 0.95 quantiles of shape variance along PC1 (right) in (A) and 0.05 quantiles of shape variance along PC1 (left) in (A). Landmarks and semilandmarks on the quadrate models are coloured according to log10-transformed per-landmark Procrustes variance. Bird species are labelled by main dietary category (see Material & Methods). Convex hulls represent the approximate area of the morphospace in A and B occupied by Galloanserae (dark gray), Passeriformes (light gray) and Psittaciformes (dashed grey line). Individual taxa and family-level clades are labelled.

PC2 captures shape variance associated with muscle attachments and articular surfaces (Figs. 3 and S16, PC2). Taxa with curved quadrate bodies, curved lateral and medial crests, and widely spaced capitula on the otic process exhibit positive scores along PC2 (Figs. 3 and S16, PC2 positive). Such quadrates also exhibit a lateromedially curved articular surface of the squamosal capitulum with an ovoid outline and a medially facing otic capitulum due to a markedly curved medial crest (best typified in Strigiformes; Fig. 3A; Fig. S13 A, E-G). These quadrate shapes are also characterised by relative short and robust orbital processes pointing lateroventrally, and ventrally facing quadratojugal cotylae with relatively small articular fossae (Fig. S16, PC2 positive), as seen in the Imperial Cormorant (*Leucocarbo atriceps*) and nightjars (Caprimulgidae), and mandibular processes with dorsally tilted caudal condyles divided by a deep groove (intercondylar sulcus; Fig. 3A; Fig. S13 A, E-G). By contrast, quadrates exhibiting negative PC2 scores are associated with narrowly spaced capitula on the otic process (Figs. 3A and S16, PC2 negative), as in some Anseriformes (Fig. 3A; Fig. S13 A, E-G), and a squamosal capitulum with a relatively flat, rounded articular surface (Fig. S16, PC2 negative). Such quadrates also have dorsomedially facing orbital processes with pointed tips and high aspect ratios, and mandibular processes in which the intercondylar vallecula and the caudal condyles are weakly developed, as in some parrots (Psittaciformes; dashed line in Fig. 3) or the Hoatzin (*Opisthocomus hoazin*; Fig. 3A; Fig. S13 A, E-G).

PC3 corresponds to shape differences in the mandibular process between parrots and all other avian lineages (Fig. 3B; Fig. S13, B, E, H-I). The parrot quadrate is mainly characterised by: 1) a mandibular process adjacent to the pterygoid condyle and significantly separated from the quadratojugal cotyle; 2) a short but robust orbital process with a blunt tip; 3) a rostrally displaced medial condyle (or rostral portion of the mandibular process); 4) a dorsally displaced caudal condyle (or caudal portion of the mandibular process); and 5) a caudomedially displaced lateral condyle (or lateral part of the mandibular process), which places the three main condyles in rostrocaudal alignment (Figs. 3B and S16, PC3 positive). This configuration differs from the more common condition in which quadrates bear a longer orbital process with a pointed tip and the mandibular process is widely separated from the pterygoid condyle but close to the quadratojugal cotyle, with three clear condyles forming a triangle or a L-shape in ventral view, such as Passeriformes (Fig. 3B; Fig.S16, PC3 negative). Shape changes associated with the remaining main axes of shape variation are detailed in the Supplementary Materials.

### (b) Quadrate shape and species autecology

Our phylogenetic generalized least squares (PGLS) regressions illustrate that all ecological factors investigated (i.e. trophic level, diet, use of beak during feeding, primary habitat, and primary lifestyle) exhibit non-significant correlations with quadrate shape (Table 1), including when we corrected for the effects of other potential explanatory factors such as the main effects of phylogenetic group and allometry, and group-specific allometries (Table S2). Similar quadrate shapes are associated with widely varying dietary ecologies (i.e. one-to-many mapping of form to ecology [8]) and very different quadrate shapes are associated with similar dietary ecologies (i.e. many-to-one mapping of form to ecology; Fig. 4).

**Fig. 4.**
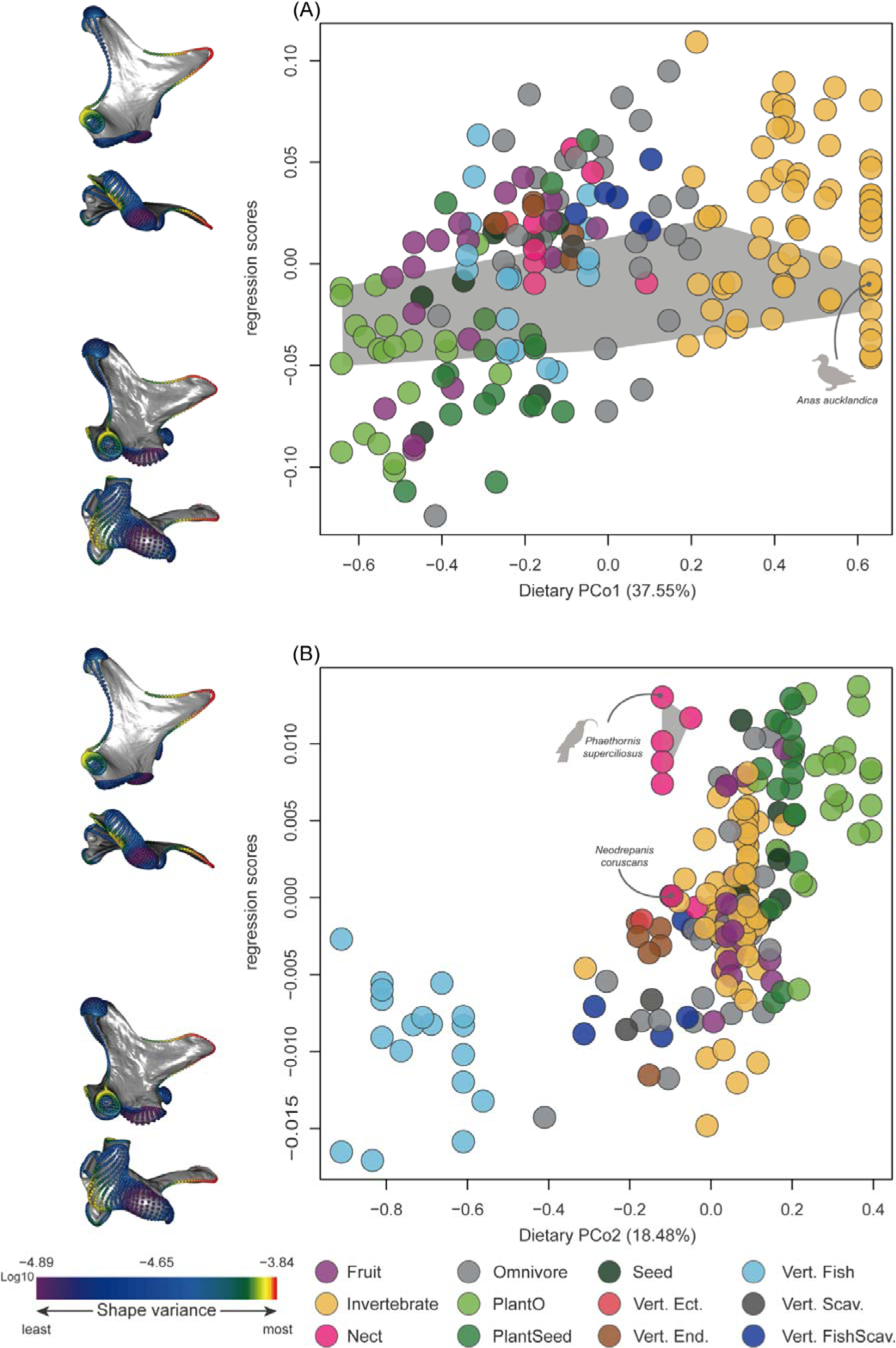
The evolutionary relationship between quadrate shape and the two main axes of dietary variation in Neornithes. Relationship between quadrate shape (regression scores [shape variance∼ Diet matrix]) and scores from the first two PCo axes (PCo1(A) and PCo2(B)) of semiquantitative dietary variation (see Methods). Bird species are labelled by their main dietary category (see Methods). Quadrate models represent the 0.95 and 0.05 quantiles of shape variance along the regression vector that is maximally associated with each of the major axes of shape variation. Quadrates are shown in lateral and ventral views. Landmarks and semilandmarks on the quadrate models are coloured by log10-transformed per-landmark Procrustes variance. Shaded area in A indicates the variance exhibited by Anseriformes; shaded area in B indicates the variance displayed by Trochilidae.

**Table 1.**
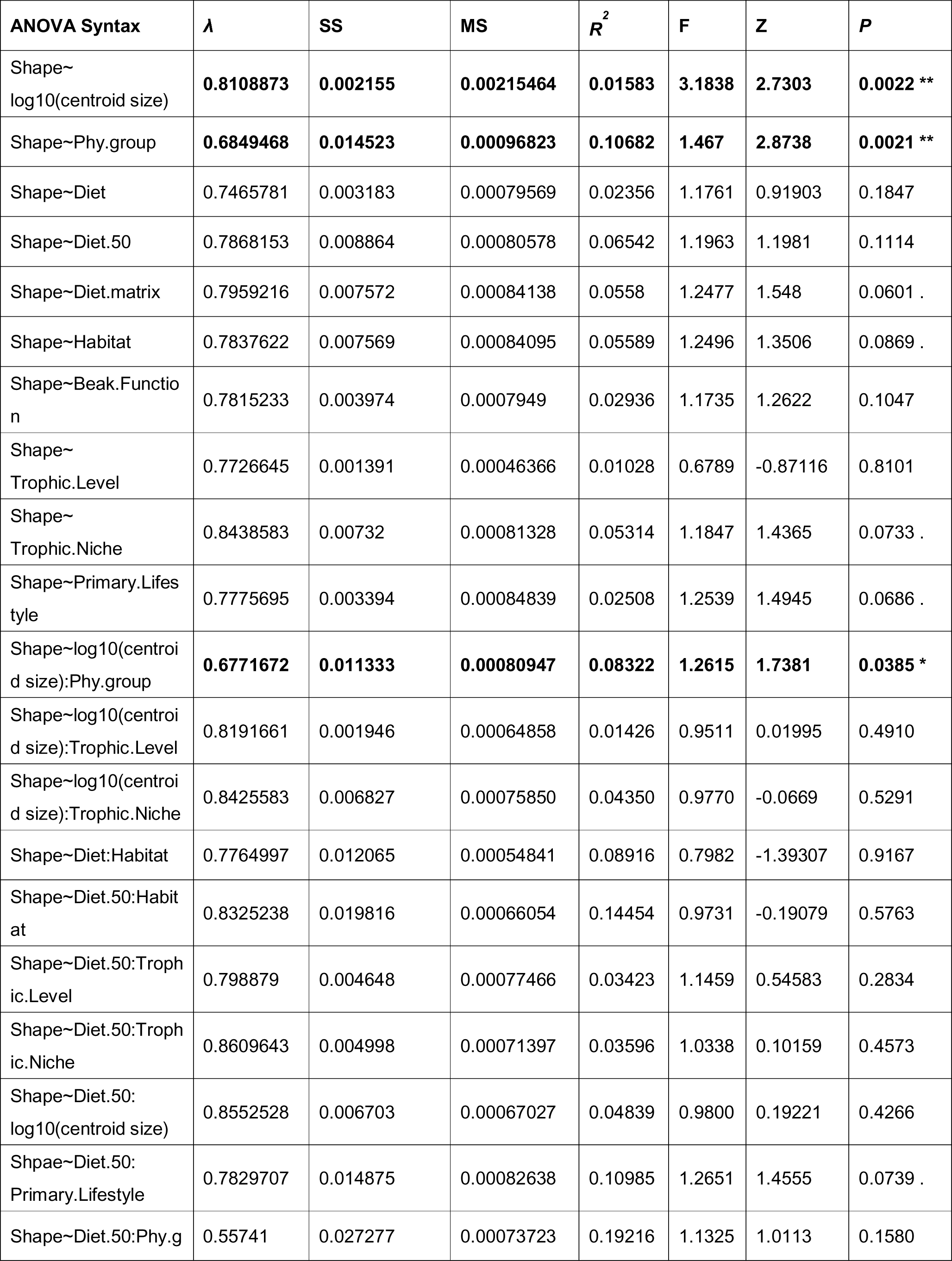
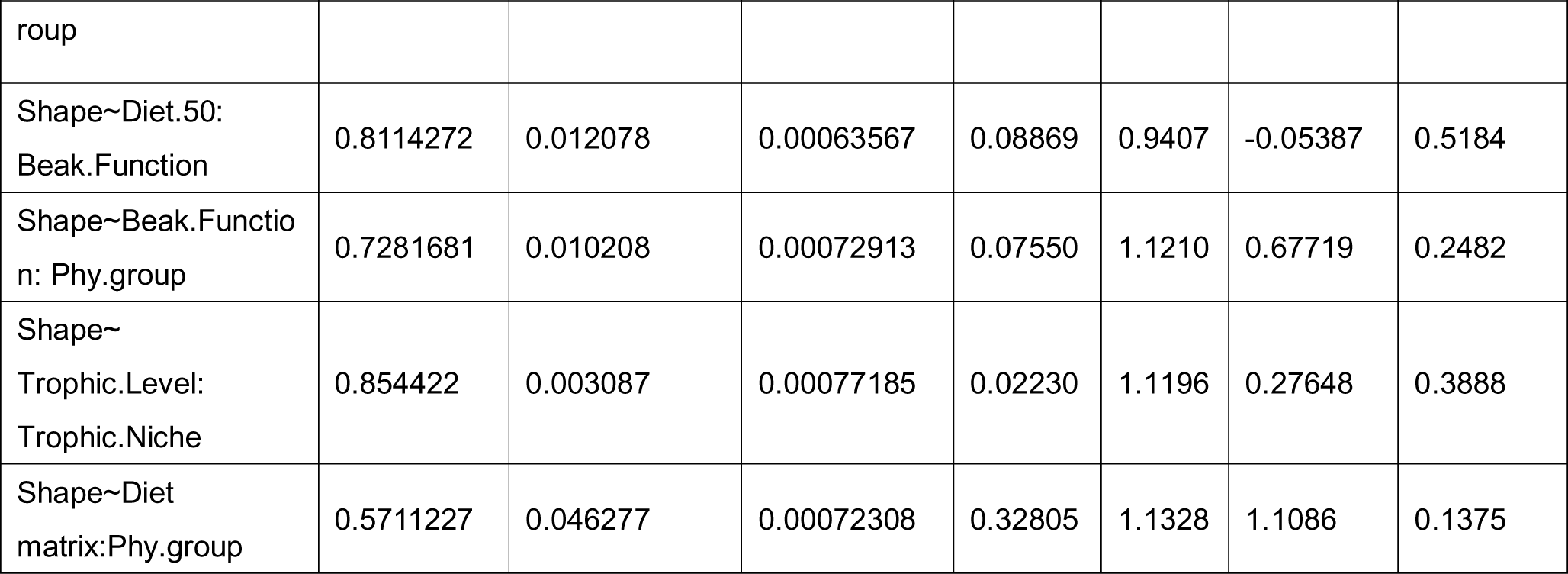
Summary of the PGLS liner models for Procrustes coordinates (quadrate shape) in Neornithes (n = 200) as a function of ecology information and its interaction. Numbers in bold indicate statistical significance (*p* < 0.05).

### (c) Quadrate shape, avian phylogeny and allometry

The strongest correlation found across Neornithes is between quadrate shape and phylogenetic group with *R^2^* of ∼0.11, meaning that roughly eleven percent of total quadrate shape variation can be explained by the major phylogenetic lineage that the quadrate-bearer belongs to (Table 1 and Fig. S24 A). This comparatively significant relationship between clade and quadrate shape can be observed in the general distribution of clades in quadrate morphospace (Fig. S13) and in the paradigmatic case of galloanserans that retain a similar quadrate morphology despite great variation in their feeding ecology (shaded area in Fig. 4A). Furthermore, we found a significant correlation between quadrate shape and size (quadrate allometry) which also varies depending on phylogenetic group (significant interaction log_10_ (centroid size):phylogenetic group in Table 1 and Fig. S24 B). This is also the case for non-Telluraves (non-landbirds) which represent most of the species in our dataset (n=137) with *R^2^* of ∼0.02 (Table S4 and Fig. S24 C), but this relationship is non-significant for Telluraves (Table S3; N = 63).

### (d) Quadrate shape, skull shape and internal integration

Quadrate shape strongly covaries with cranial (*R^2^* = 0.6525, Z = 4.25602, *p* = 1e-04) and mandibular shape (*R^2^* = 0.5395, Z = 1.97832, *p* = 0.0249), but also with the shape of the rostral portion of the cranium (*R^2^* = 0.6389, Z = 4.04526, *p* = 1e-04) and the palatine-pterygoid complex (*R^2^* = 0.5911, Z = 3.22512, *p* = 5e-04) (PPC, mostly representing morphological variance of the palatine, see [46]; Fig. 5A, Table S6). The shapes of individual regions of the quadrate also exhibit strong covariation with corresponding articular skeletal elements in the cranium and mandible; for instance, the otic process and the mandible (*R^2^* = 0.4396, Z = 2.0986, *p* = 0.0191), the otic process and the whole cranium (*R^2^* = 0.508, Z = 3.24455, *p* = 4e-04), the quadrate body and the whole cranium (*R^2^* = 0.6458, Z = 4.7964, *p* = 1e-04), the quadratojugal cotyle and the mandible (*R^2^* = 0.5327, Z = 3.38037, *p* = 1e-04), the quadratojugal cotyle and the jugal bar (*R^2^* = 0.3952, Z = 2.39022, *p* = 0.0073), and the quadratojugal cotyle and the rostrum (*R^2^* = 0.369, Z = 2.07642, *p* = 0.019). These relationships are generally invariant when analysed at lower phylogenetic scales within both Telluraves and non-Telluraves (Fig. 5B-C; Tables S7 and S8). For instance, within Telluraves (n = 40), we found numerous strong relationships between the quadrate bone and adjacent skeletal elements, such as the quadrate and the mandible (*R^2^* = 0.8242, Z = 3.06459, *p* = 2e-04), the quadrate and the whole cranium (*R^2^* = 0.746, Z = 2.4804, *p* = 0.0043), and the quadrate and the PPC (*R^2^* = 0.7878, Z = 2.95048, *p* = 6e-04; see full list of significant associations in Table S7). Within non-Telluraves (n = 57), we found strong integration between the quadrate and the mandible (*R^2^* = 0.6768, Z = 2.55903, *p* = 0.0044), the quadrate and the cranium (*R^2^* = 0.6959, Z = 3.07433, *p* = 0.001), and the quadrate and the PPC (*R^2^* = 0.6558, Z = 2.46721, *p* = 0.0064; see full list of significant association in Table S8). Significant shape covariation (*p* value > 0.05) between specific processes and adjacent bones of the cranium and mandible reveal a more complex pattern than entire quadrate shapes, and inferred patterns of shape integration differ slightly among taxonomic subsets. For instance, Telluraves exhibits a different covariation structure than Neornithes and non-Telluraves in terms of the relationship between the orbital portion of the quadrate (orbital process + basiorbital fossa) and mandible, the otic process and cranium, the quadrate body and mandible, the pterygoid condyle and PPC, and the pterygoid condyle and mandible (Tables S6-S8, Mandibular process vs. Mandible). Significant associations between the orbital portion of the quadrate and mandible, pterygoid condyle and mandible, and pterygoid condyle and PPC are only observed in Telluraves. This stronger covariance is potentially related to specific muscle attachments or patterns of force transmission within the palate system that are restricted to this major neornithine subclade. Meanwhile, strong relationships between the geometry of the otic process and cranium are generally observed across crown birds and among non-telluravians. Shape covariation among all quadrate subregions is uniformly statistically significant and stronger than covariation between quadrate shape and other regions of the skull (Fig. 5; Table S6).

**Fig. 5.**
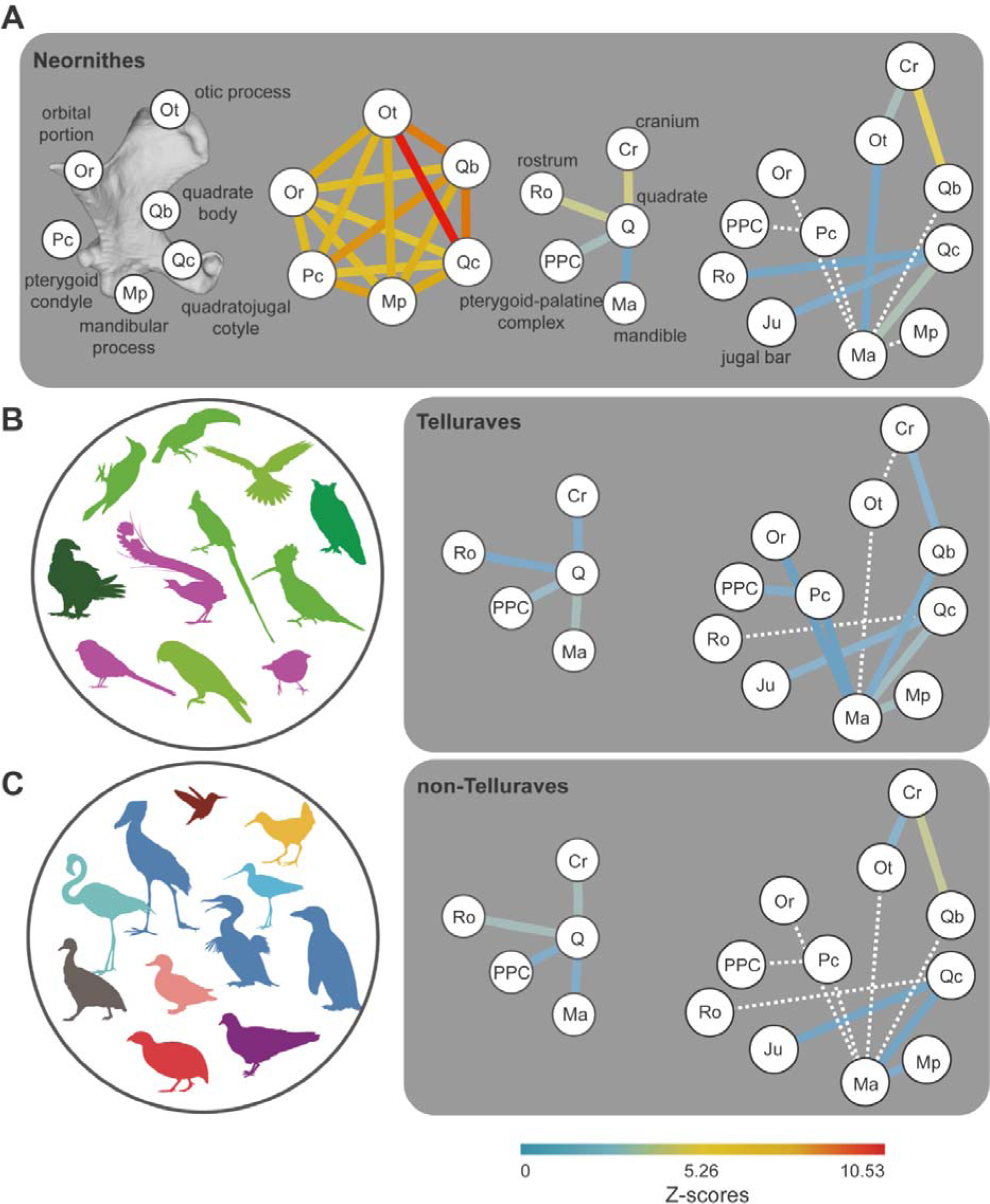
Evolutionary integration of the quadrate in Neornithes, Telluraves and non-Telluraves Neornithes. (A) Integration between each area/process of the quadrate in Neornithes (n=200), between the quadrate and its neighbouring bones and between different areas/processes of the quadrate and its neighbouring bones in Neornithes (n=97). (B) Integration between the quadrate and its neighbouring bones and between different areas/processes of the quadrate and its neighbouring bones in Telluraves (n=40). (C) Integration between the quadrate and its neighbouring bones and between different areas/processes of the quadrate and its neighbouring bones in non-Telluraves (n=57). Dashed lines indicate non-significant relationships.

## 4. Discussion

The quadrate is a key component of the kinetic system of the skull in neornithine birds (Fig. 1) [18–21], and its functional relevance implies that strong selective pressures directly related to feeding and foraging ecology may have shaped its morphological evolution. However, our results challenge this assumption: the relationship between quadrate shape and feeding and foraging autecology in extant birds is generally either non-significant or weak (although variable, see below) at multiple phylogenetic scales (Tables 1 and S3 and S4), even when accounting for potential confounding factors (Table S2). As such, the diversification of quadrate shape in Neornithes may not have been driven directly by selection for ecological optimality. Many-to-one (different shapes associated with the same ecology) and one-to-many (similar shapes associated with disparate ecologies) relationships between quadrate morphology and function are apparent from our results (Figs. 3 & 4). For instance, nectarivory, a highly specialised feeding ecology, is represented in our dataset by several distantly related species which all exhibit highly divergent quadrate shapes (i.e. many-to-one relationship of morphology and function; Fig. 4B), perhaps reflecting different mechanisms of nectar feeding associated with distinct phylogenetic histories [47], and/or that the morphology and function of other areas of the head like the tongue (including the hyoid skeleton) and/or the upper and lower jaws are simply more closely linked to nectar feeding than the quadrate is [48]. These results echo recent studies that have demonstrated that, at broad phylogenetic scales, avian evolution is characterised by complex relationships between skeletal morphology and ecology. For instance, dietary or other bill usage traits explain a relatively small portion of cranial shape variance across Neornithes [8–9], despite the fact that these are among the strongest form-function relationships across the avian skeleton [10].

We found a much stronger, statistically significant link between avian quadrate geometry and phylogeny, with different avian lineages exhibiting distinctive quadrate shapes and clade-specific allometries (see Fig. 4; relevant tables). For instance, Anatidae (ducks, geese, swans) exhibit a wide spectrum of feeding ecologies ranging from piscivory (e.g., *Mergus*) to herbivory (e.g., *Sarkidiornis*), with beak morphologies reflecting this ecological disparity [11]. However, despite this dietary disparity, anatids exhibit very similar quadrate morphologies overall (i.e. a one-to-many relationship of morphology to function; Fig. 4A, S17, and Table S5). That said, one of our ecological categories (trophic level) strongly correlates with the limited quadrate shape variance observed in Anatidae (Table S5), indicating that shape may link to functional characters or autecology on more restricted phylogenetic scales (that is, within, but perhaps not among, major clades). Certain notable morphological features of the quadrate represent autapomorphies of specific avian subclades, such as the caudomedial extension of the otic capitulum owls (Strigiformes), which effectively creates two distantly spaced capitula on the otic process (character 548 in [50]). The orbital process in nightjars (Caprimulgidae) is weakly developed with a low aspect ratio, related to the loss of the *m. pseudotemporalis profundus* [51]. However, the reduction of the orbital process enables a greater gape size in nightjars [51] and is related to the reorientation of the lower jaw when closed [52]. The mandibular process of Psittaciformes (parrots) is characterized by a single, deep medial condyle with a prominent articular surface, which enables the quadrate to slide rostrocaudally along its articular surface with the lower jaw to facilitate strong adduction [53]. The relatively strong correspondence we find between quadrate shape and phylogeny is in line with these examples, with distinctive autapomorphies characterising several important avian subclades, highlighting the quadrate as a potential source of useful phylogenetic characters in diagnoses of fossil birds [50, 54–56].

Our study focused on the evolutionary relationships between quadrate shape and ecology; however, this may conceal informative relationships between function and other aspects of quadrate form, such as size. The size of morphological structures has been shown to predict important aspects of avian ecology (e.g., sternum [57–58], cranium and mandible [10]; and labyrinth [59–60]), and it stands to reason that relative quadrate dimensions may provide important, though as-yet unexplored insights into cranial function. The important role of the quadrate in avian cranial kinesis relates to its contribution to a mechanical four-bar linkage that drives elevation of the upper beak with respect to the cranium [61]. Although size variation (centroid size) of the avian quadrate is only weakly associated with shape variance (∼ 1%, Table 1), the relative size of the quadrate (with respect to the other cranial elements), rather than geometric shape, may prove to be a more important aspect of quadrate morphology due to its influence on mechanical advantage of the suspensorium and palate, a question that should be investigated in the future.

Our results investigating bird quadrates are strikingly different to those obtained in a recent study of squamate quadrates, which showed that ecological habits explain a relatively large proportion of quadrate shape variance (20%), with little shape variance explained by phylogeny [62]. These divergent results in birds and squamates may have multiple explanations. For instance, there is some evidence that the bird quadrate may exhibit a comparatively stronger degree of evolutionary integration with the palatine bones than the squamate quadrate [63–64] such that quadrate form in birds may be dictated to a greater extent by complex patterns of cranial structural integration rather than by selection for mechanical optimality for specific foraging modalities acting on individual cranial elements. Indeed, our analyses support this view: we found evidence for strong evolutionary integration between the shape of the quadrate and that of adjacent bony elements of the skull, such as the palate, at all phylogenetic scales (Neornithes, Telluraves, and non-Telluraves), indicating that morphological evolution of the avian quadrate has been shaped to some degree by the evolution of the adjacent head skeleton. Nonetheless, our results also support the hypothesis that the quadrate has evolved as a coherent unit (i.e. an evolutionary module) within the avian cranial kinetic system as indicated by a stronger degree of covariation among regions within this element than between the quadrate and adjacent skeletal components (Fig. 5).

The integration between quadrate form and adjacent skeletal elements most likely reflects functional associations between components of the avian kinetic system. The nature of these relationships, however, may be lineage specific. For instance, in landbirds (Telluraves), the high degree of shape covariation between the quadrate body and the mandible (Z = 2.4826), and between the pterygoid condyle of the quadrate and the PPC (pterygoid + palatine; Z = 2.48699) may reflect a higher degree of functional integration of these cranial elements in this clade than in other avian lineages, with potential functional consequences such as greater efficiency of force transmission from the back of the skull to the tip of the rostrum (Fig. 5C and Table S6). Furthermore, the mandibular process of the quadrate does not significantly covary in shape with the shape of the mandible across Neornithes as a whole, but these bones covary strongly when Telluraves (Z = 3.14419) and non-Telluraves (Z = 2.34769) are analysed separately. These variable relationships may reflect functional differences in the kinetic systems of landbirds with respect to other major neornithine lineages. Another example is the greater degree of integration between the quadrate’s orbital portion (orbital process + basiorbital fossa) and the mandible (Z = 1.72511) seen only in Telluraves, which may reflect the influence of the attachment of *M. pseudotemporalis profundus*, a key muscle in the kinetic system, as the extent of its development is proportional to that of the orbital process [65] (Fig. 5C and Table S6).

## Conclusions

Collectively, our results underline the complexity of the evolution of the avian feeding apparatus and suggest that different lineages have frequently arrived at functionally comparable, yet morphologically divergent solutions. The three-dimensional complexity of avian quadrate geometry and the quadrate’s tight functional integration within the suspensorium/palate region may underlie the weak relationship between quadrate geometry and feeding autecology at a broad macroevolutionary scale, which appears to be far weaker than the relationships emerging from studies of craniofacial ecomorphology [e.g., 8-9]. Meanwhile, although our results suggest that quadrate shape variance is largely decoupled from feeding autecology or patterns of beak usage at a broad macroevolutionary scale, the strongly integrated relationship between quadrate geometry (including the shape of its articular surfaces and muscle attachments) and adjacent bones within the skull suggest shape-related functional differences within the cranial kinetic system of different groups of birds. Future studies explicitly interrogating the biomechanical role of the quadrate within the avian cranial kinetic system should focus on additional aspects of quadrate morphometry (e.g., the width of the otic process and the quadrate body, the relative size of the orbital process to the quadrate body, or the width of the mandibular process) to yield a deeper understanding of the relationship between quadrate morphology and beak function across extant bird phylogeny. Importantly, the significant association between quadrate shape and phylogeny highlights opportunities to incorporate quadrate geometry into investigations aiming to place fossil bird remains in a phylogenetic context using an explicitly quantitative framework. The frequency of well-preserved quadrates in the avian fossil record highlights the potential of this line of research for shedding light on the early evolutionary history of crown birds [e.g., 66-68].

## Acknowledgments

We thank K. Smithson (Cambridge Biotomography Centre) for access to micro-CT scanning facilities. We thank P. Smith (Oxford University Museum of Natural History) for providing the micro-CT scanning images of Dodo (*Raphus cucullatus*, OUMNH:ZC:11605). This work was funded by UKRI grant MR/X015130/1 to D. J. F. Additional funding for the project was provided by the European Research Council Starting Grant: TEMPO (ERC-2015-STG-677774) to R. B. J. B. For the purpose of open access, the authors have applied a Creative Commons Attribution (CC BY) licence to any Author Accepted Manuscript version arising.

